# *Ex vivo* human brain volumetry: validation of magnetic resonance imaging measurements

**DOI:** 10.1101/2025.02.27.640654

**Authors:** Amy Gérin-Lajoie, Walter Adame-Gonzalez, Eve-Marie Frigon, Liana Guerra Sanches, Anna Nayouf, Denis Boire, Mahsa Dadar, Josefina Maranzano

**Affiliations:** Department of Anatomy, Université du Québec à Trois-Rivières, Trois-Rivières, Québec, Canada; Department of Psychiatry, McGill University, Montreal, Quebec, Canada; Douglas Mental Health University Institute, Montreal, Quebec, Canada; Department of Neurology and Neurosurgery, Montreal Neurological Institute, McGill University, Montreal, Quebec, Canada

**Keywords:** MRI, segmentation, volumetry, *ex vivo*, water displacement

## Abstract

**Background:** Neurodegenerative diseases are associated with brain atrophy. The volume of *in vivo* human brains is determined with various magnetic resonance imaging (MRI) measurement tools of which the validity has not been assessed against a gold standard. Here, we propose to validate the MRI brain volumes by scanning *ex vivo-in situ* specimens (i.e., anatomical heads), which allows the extraction of the brain after the scan to compare its volume with the gold standard water displacement method (WDM).

**Methods:** We acquired 3T MRI T2-weighted, T1-weighted, and MP2RAGE images of seven anatomical heads fixed with an alcohol-formaldehyde solution routinely used in anatomy laboratories and segmented the gray and white matter of the brain using two methods: 1) a manual intensity-based threshold segmentation using Display (MINC-ToolKit, McConnell BIC), and 2) an automatic Deep-Learning-based segmentation tool (SynthSeg). The brains were then extracted, and their volumes were measured with the WDM after the removal of their meninges and a midsagittal cut (to allow water penetration into the ventricles). Volumes from all methods were compared to the ground truth (WDM volumes) using a repeated-measures ANOVA.

**Results:** Mean brain volumes, in cubic centimeters, were 1111.14±121.78 for WDM, 1020.29±70.01 for manual T2-weighted, 1056.29±90.54 for automatic T2-weighted, 1094.69±100.51 for automatic T1-weighted, 1066.56±96.52 for automatic MP2RAGE INV1, and 1156.18±121.87 for MP2RAGE INV2. All volumetry methods were significantly different (F=17.874; p<0.001) from the WDM volumes, except the automatic T1-weighted volumes.

**Conclusion:** We demonstrate that SynthSeg accurately determines the brain volume in *ex vivo-in situ* T1-weighted MRI scans. Our results also suggest that given the contrast similarity between our *ex vivo* and *in vivo* sequences, the brain volumes of clinical studies are most probably sufficiently accurate, with some degree of underestimation depending on the sequence used.

## 1. Introduction

Magnetic resonance imaging (MRI) can be used to generate three dimensional (3D) images of the human brain. From these images, scientists can observe and/or measure the contrast between gray matter (GM), white matter (WM), and cerebrospinal fluid (CSF), and then perform various subsequent analyses, using image segmentation.

The segmentation of MRI scans is a widely used technique allowing the division of an image in separate homogeneous regions with similar attributes (i.e., voxel intensity, texture, and/or depth). For brain images, it is common to apply segmentation techniques to separate the three principal tissue types (i.e., GM, WM, and CSF). This can be used to measure or visualise various brain structures and/or regions, delineate lesions, analyze anatomical structures, and study pathological brain regions [1].

Two different segmentation methods have generally been applied: 1) manual methods and 2) automatic methods. The manual image segmentation method is better to control errors related to imaging artefacts and quality (e.g., subject motion artefacts can give false contrast to certain regions that could erroneously be segmented as GM by an automatic software) [1]. However, manual segmentations may also be susceptible to the evaluator subjectivity bias (e.g., inclusion of GM-WM partial volume voxels systematically as GM), and it requires more time and rater’s expertise in neuroanatomy to be accurately performed [1–5]. Conversely, automatic image segmentation requires less processing time, reduces the evaluator subjectivity bias and improves the reproducibility of analysis [3]. However, it also comes with certain limitations, such as methodological (e.g., scan parameters, MRI scanner upgrades, etc.) and physiological factors (the presence of pathological changes may affect the accuracy of automatic tools) [4, 6]. The variability of both segmentation methods can be increased when applied to scans acquired during different MRI sessions (e.g., subject motion is not the same between MRI acquisitions taken at different times), across subjects or with different scanners [7].

The results of these segmentations on 3D MR images are frequently used to measure the volume of brain structures, regions, and/or the whole brain, which is useful in a clinical setting. For instance, volumetric measures are taken on *in vivo* human brains [3] to assess the atrophy associated with various pathologies, such as neurodegenerative diseases (e.g., Alzheimer’s disease) and multiple sclerosis [8, 9]. The validation of volumetric measures is important to ensure the accuracy and consistency of the obtained volume and the resulting diagnostic interpretation, especially in cross-sectional studies, where there are no other references for the same specimen (i.e., the measure is taken at a single point in time).

However, the total human brain volume determined *in vivo* cannot be validated by the gold standard measurement of the volume of an object by water displacement as described by Archimedes’ law, which would imply the extraction and submersion of the brain in water. Since the water displacement method (WDM) cannot be used *in vivo*, we propose an alternative validation method that stays as close to validating the *in vivo* volumes as possible. Our method consists of using *ex vivo-in situ* brains, in which anatomical head specimens from our human gross anatomy laboratory are fixed by perfusion directly inside the skull. This ensures that the brain remains in a similar environment as *in vivo* (i.e., surrounded by CSF, the meninges, the vessels, and the skull) [10, 11] and it minimises the deformation that would be caused by the manipulation of the fresh unfixed brain and its vessels during immersion fixation [10, 12]. Using *ex vivo-in situ* brains allows their extraction following the MRI acquisition protocol and their subsequent measurement with the WDM.

Hence, the goal of the present study was to validate a manual and an automatic MRI segmentation volumetric method by scanning human head specimens (i.e., *ex vivo-in situ* brains), followed by brain extraction and volume quantification using the gold standard WDM. Furthermore, we evaluated the reliability and reproducibility of the volumetric segmentations in different MRI sequences at a field strength of 3T.

## 2. Materials and methods

### 2.1 Population

We used a convenience sample of seven anatomical heads (N = 7) from the body donation program of the Anatomy Laboratory at the Université du Québec à Trois-Rivières (UQTR) (Trois-Rivières, Québec, Canada). Prior to death, donors gave their consent for body donation and sharing of their medical information for anatomical teaching and/or research purposes. The study was approved by the University’s Ethics Subcommittee on Anatomical Teaching and Research. Donors were included in the study if they had no known medical history of neurodegenerative, neurologic, neurodevelopmental, or psychiatric diseases. Mean age at the time of death was 87.0 years old (standard deviation (SD): 8.0; range 76.0-96.0). The male to female ratio was 3:4. The causes of death were either of a cardiovascular, respiratory, or cancerous nature. The mean interval between death and body fixation (*post-mortem* interval; PMI) was 22.6 hours (SD: 13.5; range 6.0-48.0). The mean interval between body fixation and brain extraction (extraction interval; EI) was 52.9 weeks (SD: 6.8; range 45.5-62.0). The case number, age, sex, cause of death, PMI, and EI are reported in Table 1.

**Table 1.** Specimen data.

| ID | Age | Sex | Cause of death | PMI (hours) | EI (weeks) |
| --- | --- | --- | --- | --- | --- |
| 1 | 96 | F | Cancer | 13.25 | 45.50 |
| 2 | 91 | M | Respiratory | 20.00 | 46.00 |
| 3 | 80 | F | Cancer | 23.00 | 49.25 |
| 4 | 76 | M | Cardiovascular | 48.00 | 61.75 |
| 5 | 80 | M | Cardiovascular | 30.00 | 62.00 |
| 6 | 93 | F | Cancer | 6.00 | 54.00 |
| 7 | 93 | F | Cancer | 18.00 | 52.00 |

ID = brain specimen identification; F = female; M = male; PMI = *post-mortem* interval; EI = extraction interval

### 2.2 Fixation procedure

All anatomical heads were fixed by perfusion of the whole body with an alcohol and formaldehyde solution (AFS) developed by the Anatomy Laboratory at the UQTR for dissection and teaching purposes (i.e., to minimize the rigidity of the organs and to reduce the carcinogenic effects of the typical 10% formalin solution commonly used in brain banks). The chemical components of the fixative solution are the following: 1.42% formaldehyde, 69% ethanol, 1.88% phenol, 20.7% glycerol, 0.01% sodium acetate, and 0.13% dettol. The bodies were perfused by injecting approximately 25L of fixative through the common carotid artery with a pump (41 kPa) (Duotronic III, Hygeco International Products) following the routine procedure of anatomy laboratories [10].

### 2.3 Head preparation

All heads were prepared by an experienced anatomy professor with specialised surgical training (A.N.) and by the anatomy laboratory personnel for MRI scanning following the standard preparation guidelines established for neurosurgical simulation as mentioned by Maranzano et al. (2020) [10]. In brief, the preparation consists of the laminectomy of the cervical vertebrae C6 and C7, closure of the dural sac around the spinal cord with non-distensible cotton thread (i.e., to keep all fluids inside the ventricles and subarachnoid spaces), transection of the spinal cord distal to the closed dural sac and transection of the head at the same level. The prepared heads were stored at room temperature until the MRI scan (in the next 6 to 36 hours following preparation) in a humidified cloth with a standard water-based conservation solution composed of glycerol and phenol.

### 2.4 MRI acquisition

The heads were scanned using a 3T Siemens Prisma scanner from the Cerebral Imaging Centre of the Douglas Research Centre (Université McGill, Montréal, Québec, Canada), where the following sequences were acquired: 3D T1-weighted (T1w) and T2-weighted (T2w), and MP2RAGE. Three contrasts of MP2RAGE were used: the first and second inversion times (INV1 and INV2) and the normalisation of both (UNI) [13]. The specific acquisition parameters [7] for each sequence are detailed in Table 2. The sequences are shown in Figure 1.

**Table 2.** MRI acquisition parameters.

| Acquisition parameters | 3D T2-SPACE | 3D T1-SPACE | MP2RAGE (INV1) | MP2RAGE (INV2) |
| --- | --- | --- | --- | --- |
| TR (ms) | 2500 | 500 | 5000 | 5000 |
| TE (ms) | 160 | 8.3 | 2.72 | 2.72 |
| FA (degrees) | 120 | 120 | 4 | 5 |
| TI (ms) | - | - | 700 | 2500 |
| Voxel size (mm<sup>3</sup>) | 0.5×0.5×0.5 | 0.5×0.5×0.5 | 1×1×1 | 1×1×1 |
| Bandwidth (Hz/px) | 620 | 620 | 260 | 260 |

TR = repetition time; ms = millisecond; TE = echo time; FA = flip angle; TI = inversion time; mm<sup>3</sup> = cubic millimeter; Hz/px = Hertz/pixel

**Figure 1.**
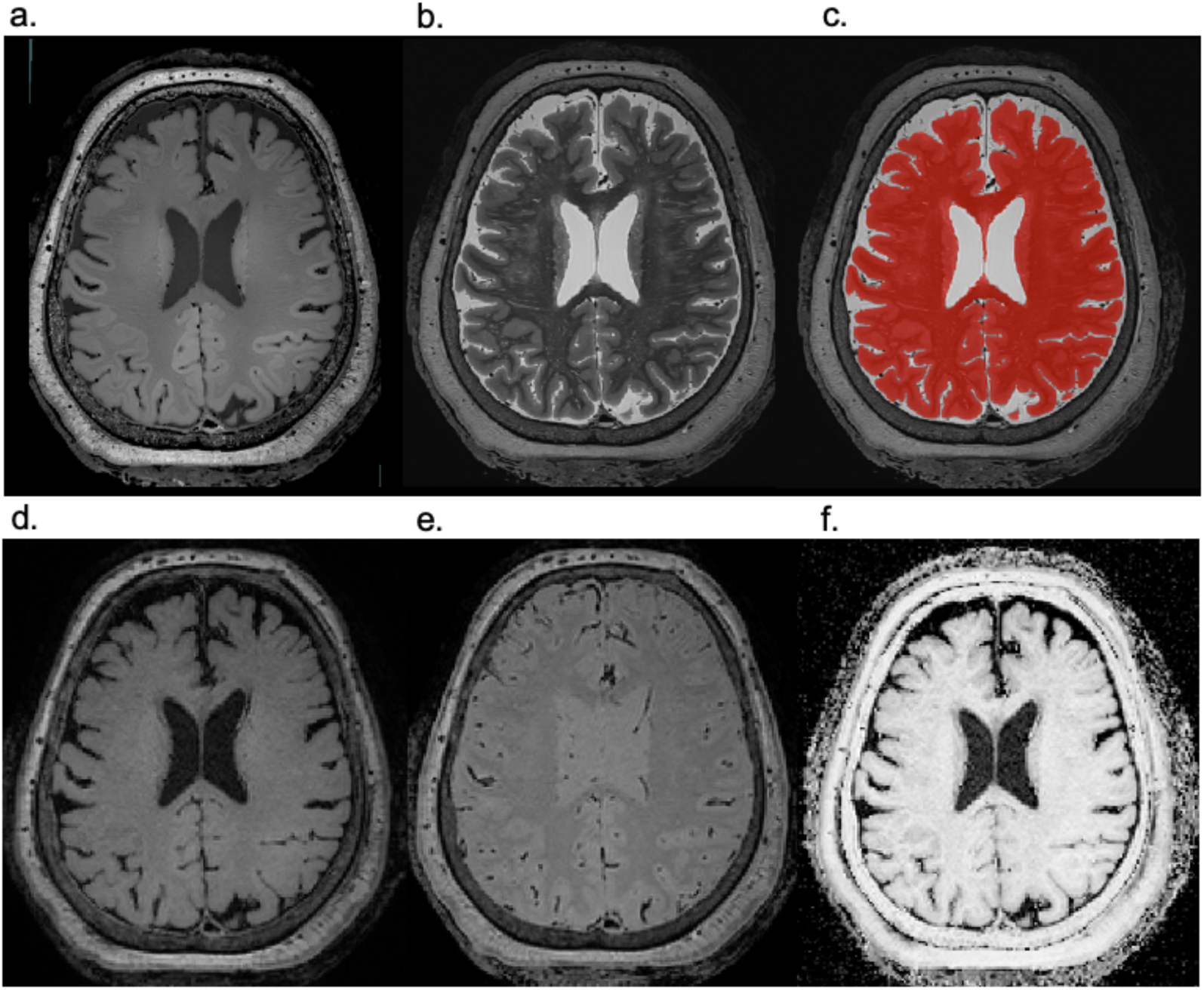
MRI segmentation volumetry. (a) shows the T1w sequence, (b) the T2w sequence, (c) the manual segmentation mask enveloping the WM and GM of the brain on the T2w sequence, (d) the MP2RAGE INV1 sequence, (e) the MP2RAGE INV2 sequence, and (f) the MP2RAGE UNI sequence. All pictures are shown in the axial axis. *WM = white matter; GM = gray matter*

### 2.5 Brain extraction

The scalp was removed with a scalpel and pliers. A circular line of opening was sawed from the frontal arcade to the external occipital protuberance, cut as low as possible to facilitate the subsequent brain extraction without damaging the brain. The dura was detached from the base of the skull with scissors. The cranial nerves, vessels, and spinal cord were sectioned to free and extract the brain, brainstem, and cerebellum from the base of the skull.

### 2.6 Water displacement volumetry

Following their extraction, we measured the volume of the brains by using the WDM based on Archimedes’ buoyancy principle, which states that an object partially or fully immersed in a liquid is buoyed up by a force that is equal to the weight of the fluid displaced by said object [14, 15]. In our case, the principle was applied by suspending the brain in a container filled with demineralised water standing on a digital scale, so that the value given by the scale (in grams) represented the weight of the water displaced by the brain itself. Knowing that the volumic mass of water is 1g = 1cm^3^, we obtained the brain’s volume by transforming the grams given by the scale in cubic centimetres (cm^3^). The set-up we used is illustrated in Figure 2. We performed the WDM following the withdrawal of the meninges (i.e., dura mater and arachnoid) and the cutting of the brain in its midsagittal axis to ensure full water penetration in the ventricular system via the lateral ventricles, since the three foramina of the fourth ventricle, which are smaller in size, might preclude the complete penetration of water or allow a more variable penetration of water. We repeated the measure three times per brain specimen (changing the water in between) to obtain a mean measure for the WDM volumes that allowed us to calculate the reliability of the WDM via an intraclass correlation coefficient.

**Figure 2.**
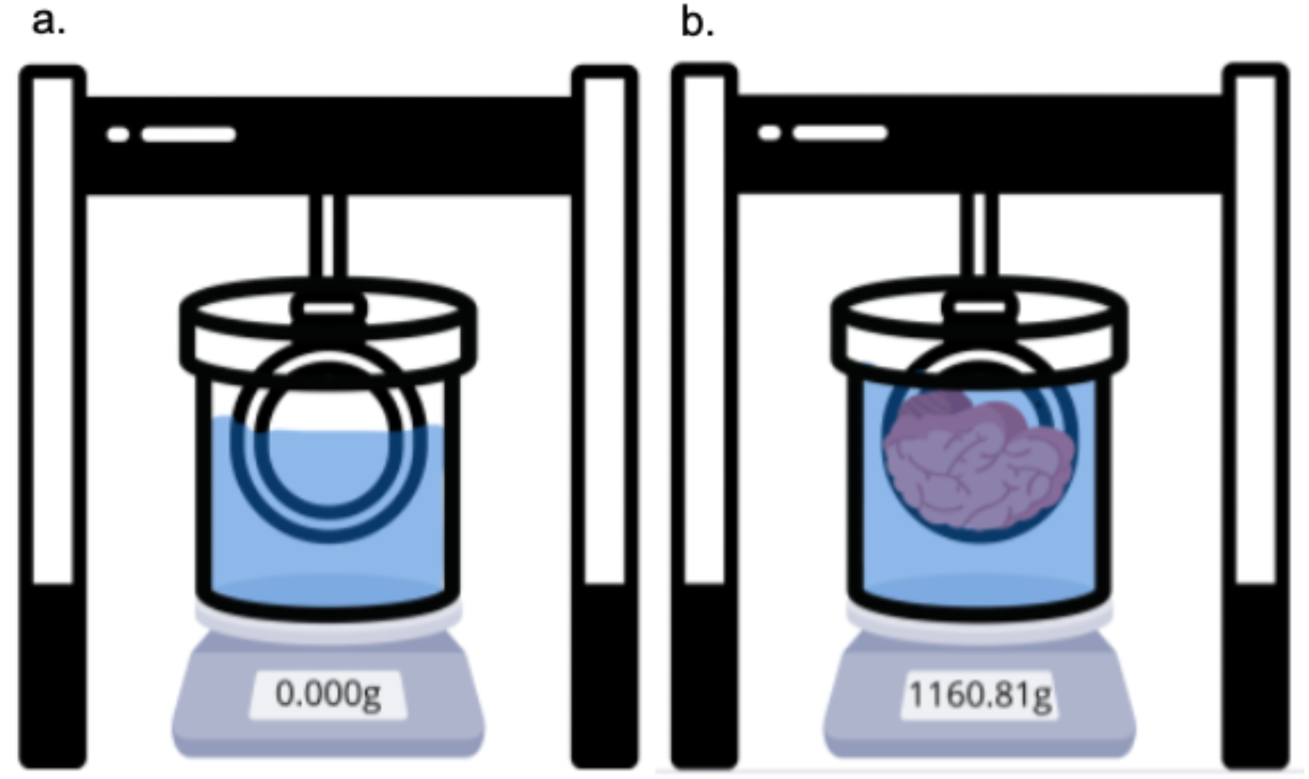
Water displacement method set-up. (a) shows the set-up before the brain is immersed in water. (b) shows the set-up with the suspended brain immersed in water. *g = gram*

### 2.7 MRI manual segmentation volumetry

We proceeded to segment the axial T2-weighted MR images with a manual method using the Display software (MINC-ToolKit, McConnell BIC). First, the software turned the DICOM format of the MR images to a MINC format so that they could be read with Display. We then created a mask for the entire brain to obtain the total volume in cubic centimeters by segmenting the WM and GM of the brain hemispheres, brainstem, and cerebellum (see Figure 1). The CSF was left out of the mask, but the choroid plexuses in the ventricles were included in the segmentation. We performed the segmentation on axial planes from top to bottom, by starting with the first images showing brain tissue at the top of the calvarium and finishing with the last images showing cerebellar tissue at the base of the skull. We used the threshold function of the Display software to segment the images with an intensity-based approach. However, since the threshold function only takes the voxel intensity into account without considering the different tissue types or the underlying anatomy, the masks were adjusted manually to delete false positive voxels (e.g., superficial vessels) and add false negative voxels (e.g., most superficial cortical GM) to the segmentation. The segmentations were first performed by an experienced rater in neuroanatomy (A.G.L., with three years of experience), then blindly repeated a second time at a later moment (i.e., more than two weeks later) to determine intra-rater variability of the manual segmentation method. The segmentations were also independently performed by two other raters with a combined experience of 29 years in neuroanatomy and brain imaging (J.M. and E.M.F.) to determine inter-rater variability of the semi-manual segmentation method.

### 2.8 MRI automatic segmentation volumetry

We proceeded to segment the axial T2-weighted MR images with an automatic method using SynthSeg [16], a Deep-Learning-based segmentation method capable of segmenting 3D brain images across multiple structural MRI modalities. SynthSeg was trained using synthetically generated images produced from segmentation masks with a wide range of inter-tissue contrasts, making it a suitable option for *post-mortem* brain image segmentation.

The files containing the MR images were converted from a MINC (*.mnc*) format to a Nifti (.*nii*) format compatible with the SynthSeg software using the *mnc2nii* function available in the MINC-ToolKit package. Since SynthSeg can only generate segmentation masks at the standard 1 mm isotropic voxel size, the inputs were resampled by the software prior to their segmentation using trilinear interpolation to match the desired standard 1 mm isotropic voxel size. Accordingly, the automatically generated segmentation masks were then rescaled to the native space (i.e., 0.5 mm isotropic voxel size) using nearest-neighbor interpolation with FreeSurfer’s *mri_vol2vol* function with the *--regheader* option enabled [17]. This allowed the computation of segmentation agreements and enabled us to compare volumetry masks between the manual and automatic labels. Finally, we merged the generated labels into a binary brain mask to match the manual binary segmentation mask generated using Display and converted the files back into a MINC format using the *nii2mnc* available in the MINC-ToolKit package.

To evaluate the MRI automatic segmentation method across different MRI sequences, we used the same procedure to segment the images of the same brain specimens in T1-weighted images, and MP2RAGE (INV1, INV2, and UNI).

### 2.9 Statistical analysis

Data from all six different volumetry methods (i.e., WDM, MRI T2w manual segmentation, MRI T2w automatic segmentation, MRI T1w automatic segmentation, MRI MP2RAGE INV1 automatic segmentation, and MRI MP2RAGE INV2 automatic segmentation) were compared using descriptive statistics (mean and SD given their normal distribution) and a repeated measures ANOVA with SPSS (version 29.0.1.1). Simple contrasts were applied to compare every MRI segmentation method with the gold standard (i.e., the WDM). A repeated measures ANOVA with a Bonferroni correction was also used to compare the automatic segmentation methods between one another. The boxplots were made using GraphPad Prism (version 10.4.1). Finally, a paired t-test was used to assess the differences between the manual and automatic segmentations on the T2w sequence.

Reliability within the MRI manual segmentation method was assessed by calculating the Dice kappa, Jaccard kappa and intraclass correlation coefficient (ICC). Both kappa coefficients were calculated with the MINC-ToolKit package, whereas the ICC and its 95% confident interval was calculated with SPSS based on a single-rating (for intra-rater reliability) or mean-rating (for inter-rater reliability), absolute-agreement and two-way mixed effects model.

## 3. Results

### 3.1 Water displacement volumetry

The mean brain volume measured with the WDM was 1111.14 cm^3^ (SD: 121.78) without meninges (i.e., after removal of dura mater and arachnoid) and with both hemispheres separated (i.e., after midsagittal cut to allow unencumbered water penetration into the ventricles). Individual data for every brain specimen are reported in Table 3. Intra-rater variability of the WDM applied to the three sets of volumetric measurements per brain specimen generated an ICC of 0.994 [0.947-0.999].

**Table 3.** Brain volumes measured by the WDM and the MRI segmentation method.

| ID | Volume (cm<sup>3</sup>) |  |  |  |  |  |
| --- | --- | --- | --- | --- | --- | --- |
|  | WDM | T2w |  | T1w | MP2RAGE<br>(INV1) | MP2RAGE<br>(INV2) |
|  |  | Manual | Automatic | Automatic | Automatic | Automatic |
| 1 | 991.000 | 931.958 | 954.798 | 972.820 | 944.182 | 1067.157 |
| 2 | 1222.000 | 1096.250 | 1147.128 | 1220.807 | 1164.028 | 1269.832 |
| 3 | 1068.000 | 1009.020 | 1031.796 | 1084.534 | 1070.176 | 1116.066 |
| 4 | 1144.000 | 1002.020 | 1059.771 | 1104.903 | 1074.856 | 1145.878 |
| 5 | 1316.000 | 1127.270 | 1207.065 | 1233.365 | 1215.164 | 1368.158 |
| 6 | 1028.000 | 957.623 | 986.948 | 1013.178 | 993.172 | 1109.452 |
| 7 | 1009.000 | 1017.900 | 1006.503 | 1033.250 | 1004.342 | 1016.733 |
| Mean | 1111.140 | 1020.292 | 1056.287 | 1094.694 | 1066.560 | 1156.182 |

ID = brain specimen identification; cm<sup>3</sup> = cubic centimeter; WDM = water displacement method;
T2w = T2-weighted; T1w = T1-weighted; INV1 = inversion time 1; INV2 = inversion time 2

### 3.2 MRI manual segmentation volumetry

The mean brain volume measured with the manual segmentation method on T2w images was 1020.29 cm^3^ (SD: 70.01) (using the measures from the first segmentation of the first rater). Individual data for every brain specimen are reported in Table 3.

Intra- and inter-rater reliability of the manual segmentation method respectively showed Dice kappas of 0.9745 and 0.9809, Jaccard kappas of 0.9915 and 0.9658, and ICCs of 0.967 and 0.978 (see Table 4).

**Table 4.** Reliability analyses for the MRI manual segmentation volumetry methods.

|  | **Intra-rater reliability** | **Inter-rater reliability** | **Manual vs automatic** |
| --- | --- | --- | --- |
|  | (rater 1 vs rater 1) | (rater 1 vs another rater) | **methods** |
| Dice kappa | 0.9745 | 0.9809 | 0.9481 |
| Jaccard kappa | 0.9915 | 0.9658 | 0.9636 |
| ICC | 0.9670 | 0.9870 | 0.9810 |

ICC = intraclass correlation coefficient

### 3.3 MRI automatic segmentation volumetry

The mean brain volumes measured with the automatic segmentation method in the four different sequences were 1056.287 cm^3^ (SD: 90.539) for T2w images, 1094.694 cm^3^ (SD: 100.511) for T1w images, 1066.560 cm^3^ (SD: 96.521) for MP2RAGE INV1, and 1156.182 cm^3^ (SD: 121.875) for MP2RAGE INV2. Individual data for every brain specimen are reported in Table 3.

We excluded the volumes generated for the MP2RAGE UNI sequence because they failed the quality control test. Large portions of brain tissue were not correctly segmented across all specimens. Misclassified voxels were observed in the background as well as in the scalp. The SynthSeg software also misclassified the brainstem as CSF, leading to its exclusion in the final volume. These recurrent problems are illustrated in Figure 3.

**Figure 3.**
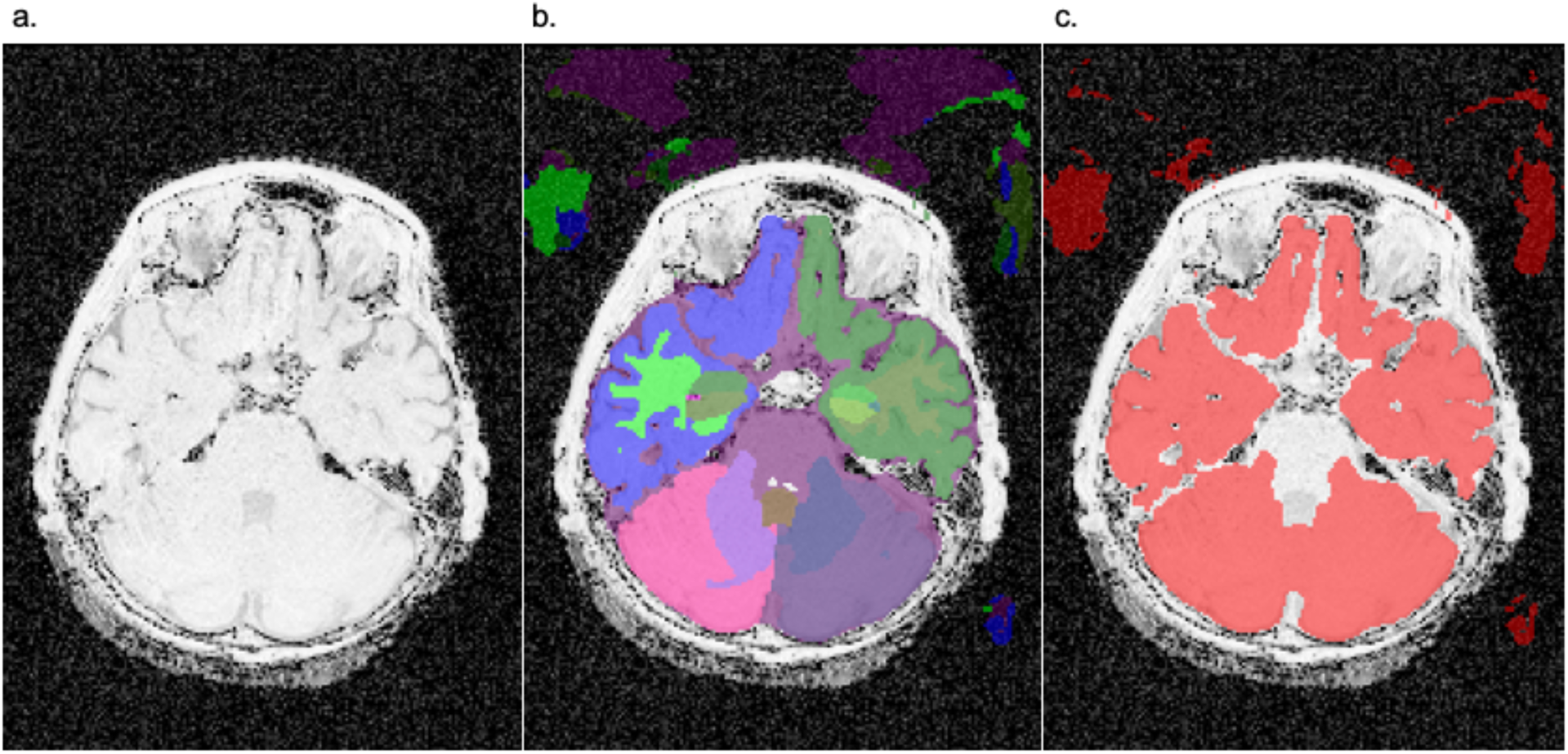
Automatic segmentation of UNI images. (a) shows the UNI image without segmentation. (b) shows the UNI image after segmentation with SynthSeg, but before merging the chosen labels. (c) shows the final segmentation mask with the chosen merged labels. There are false positive voxels in the background in both (b) and (c). The brainstem was segmented with the label for CSF (b), so it is erased during the label merging (c). This type of misclassification during the segmentation happened in all 7 cases.

### 3.4 Volumetry methods comparison

There was a significant difference across the six volumes (i.e., WDM, MRI T2w manual segmentation method, MRI T2w automatic segmentation method, MRI T1w automatic segmentation method, MRI MP2RAGE INV1 automatic segmentation method, and MRI MP2RAGE INV2 automatic segmentation method) (F_1.6;9.3_=17.874; p<0.001). Simple contrasts applied to compare every MRI segmentation method with the WDM showed that the WDM generated significantly higher volumes when compared to the MRI T2w manual segmentation (p=0.011), the T2-weighted automatic segmentation (p=0.007), and the MP2RAGE INV1 automatic segmentation (p=0.017). We also observed that the WDM generated significantly lower volumes when compared to the MRI MP2RAGE INV2 automatic segmentation (p=0.008), but there was no significant difference with the MRI T1w automatic segmentation (p=0.275). All these observations are illustrated in Figure 4. A visual representation of the location where the manual and automatic segmentation methods differ on T2w images is also shown in Figure 5.

**Figure 4.**
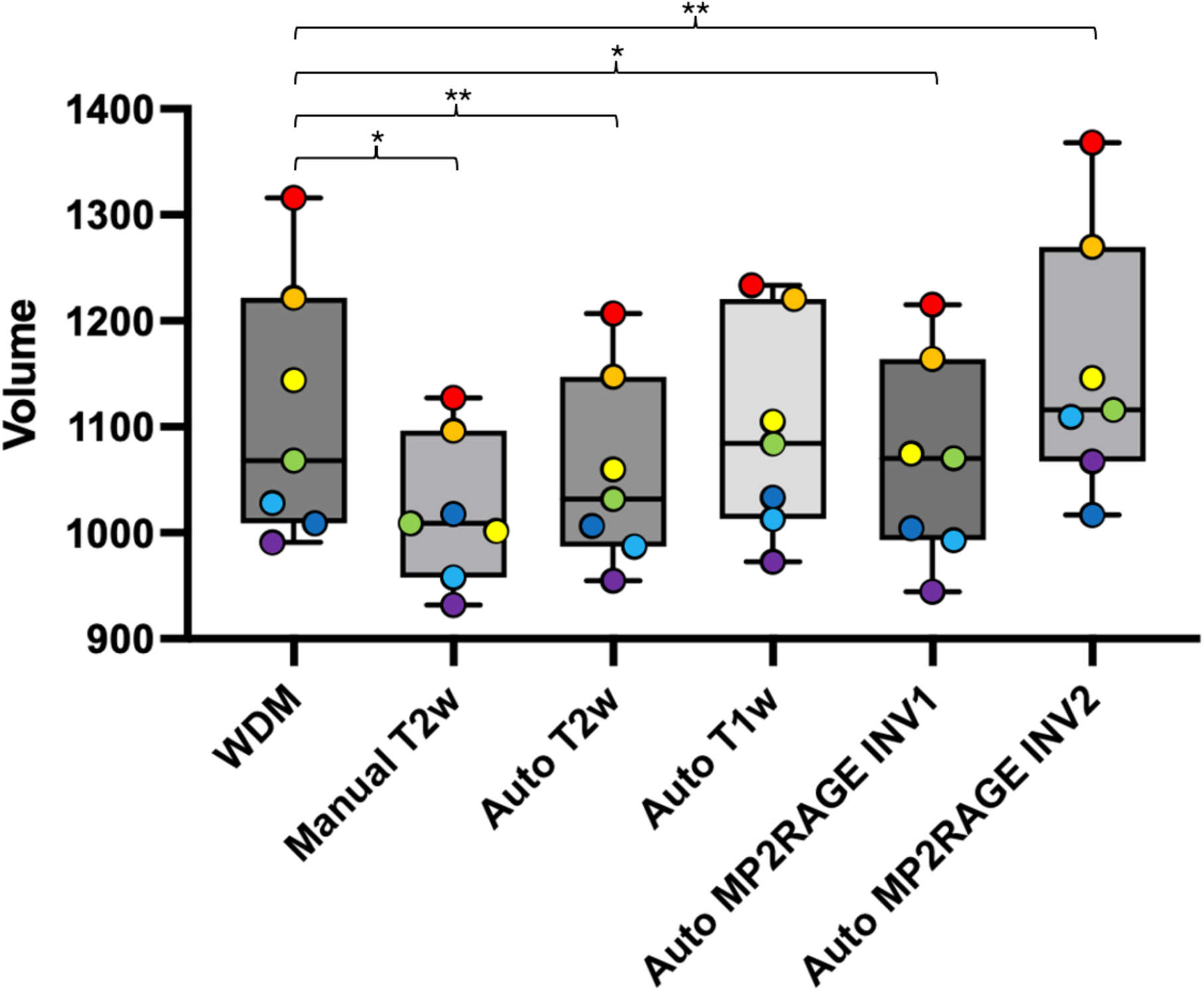
Brain volume comparisons between volumetry methods. The boxplots show the distribution of volumes measured for each of the six volumetry methods. The whiskers show the range of the distribution (minimum and maximum). The colored dots show the individual data points, and each color is associated with a specific specimen. F_1.556;9.335_=17.874; p<0.001. *\*p<0.05, **p<0.01, ***p<0.001*

**Figure 5.**
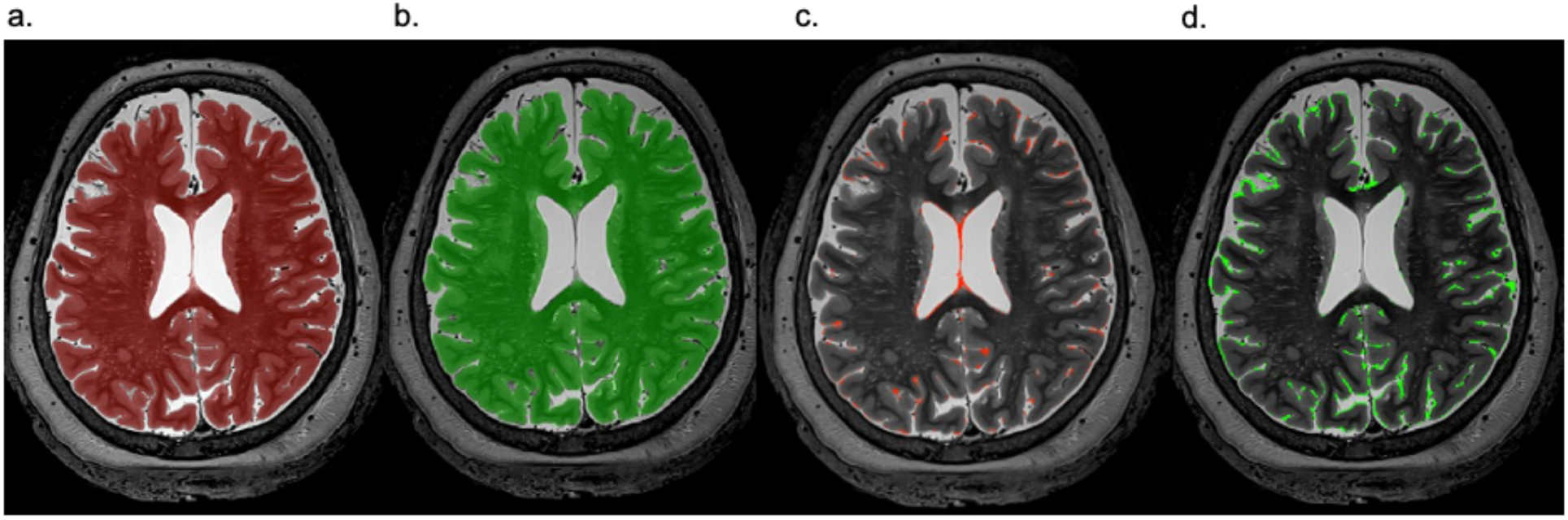
Differences between the T2w manual and automatic segmentation methods. (a) shows the MRI T2w manual segmentation mask. (b) shows the MRI T2w automatic segmentation mask. (c) shows the under-segmentation of the automatic software compared with the manual segmentation. (d) shows the over-segmentation of the automatic software compared with the manual segmentation.

We also assessed the differences across the four MRI sequences segmented with the automatic method (i.e., T2w, T1w, MP2RAGE INV1, and MP2RAGE INV2). We found that there were significant differences between the four groups (F_1.3;7.9_=21.170; p=0.001). The Bonferroni post hoc test revealed that the T2w sequence generated significantly lower volumes when compared with the T1w sequence (p=0.013) and the MP2RAGE INV2 sequence (p=0.008), but no significant difference was found with the MP2RAGE INV1 sequence (p=0.798). The test also showed that the MP2RAGE INV1 sequence generated significantly lower volumes when compared to the T1w sequence (p=0.011) and the MP2RAGE INV2 sequence (p=0.017). We found no other significant differences concerning the volumes measured by the automatic segmentation method between MRI sequences. All these observations are illustrated in Figure 6.

**Figure 6.**
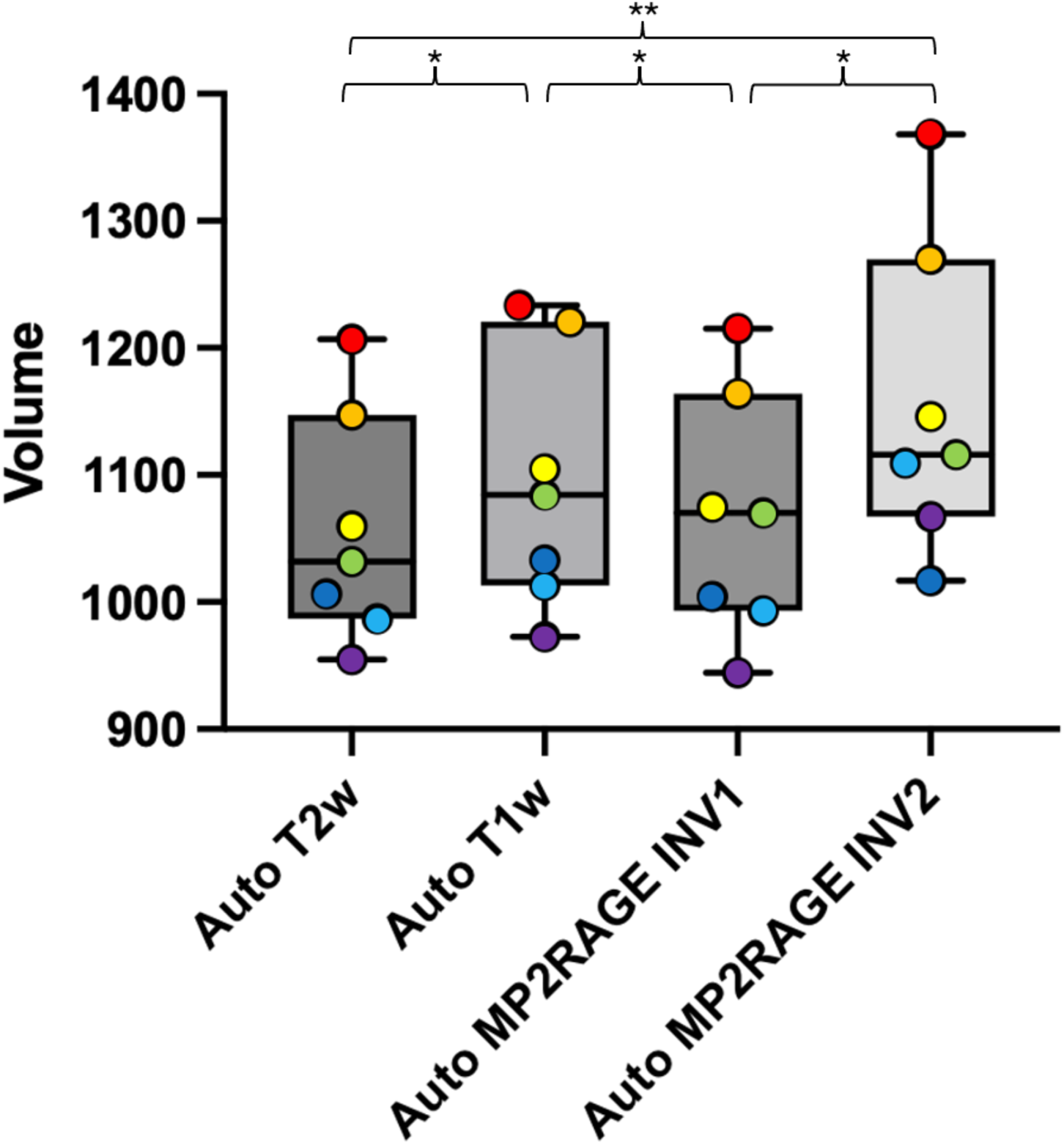
Brain volume comparisons between the automatic segmentation methods. The boxplots show the distribution of the automatically generated volumes measured for each of the four MRI sequences. The whiskers show the range of the distribution (minimum and maximum). The colored dots show the individual data points, and each color is associated with a specific specimen. F_1.3;7.9_=21.170; p=0.001. *\*p<0.05, **p<0.01, ***p<0.001*

Finally, a paired t-test showed that the brain volume in the T2w sequence was significantly lower when segmented manually as opposed to automatically (1020.29 ± 70.01 vs 1056.29 ± 90.54; *t*(6) = -3.219; p=0.018).

## 4. Discussion

In this study, we compared six volumetry methods for the *ex vivo* human brain: 1) a WDM of the brains without their meninges and separated in two hemispheres to ensure unencumbered water penetration into the ventricles, 2) a MRI T2w manual segmentation method, 3) a MRI T2w automatic segmentation method, 4) a MRI T1w automatic segmentation method, 5) a MRI MP2RAGE INV1 automatic segmentation method, and 6) a MRI MP2RAGE INV2 automatic segmentation method. Our goal was to determine the validity of both the MRI manual and automatic segmentations for measuring human brain volumes by comparing them with the gold standard method of water displacement (i.e., the WDM). Our results demonstrated a statistical difference between MRI segmentation methods and the WDM, except for the automatic segmentation generated on T1w images.

### 4.1 Water displacement volumetry

Water displacement volumetry is usually considered to be the gold standard in terms of volumetric measurements [18]. As such, we decided to use the WDM as a gold standard since it enabled us to measure the actual volume of the full brain by adapting the buoyancy principle of Archimedes, which is based on the equivalence between the weight of the water displaced by an immersed object (i.e., the brain) into cubic centimetres [14, 15]. This technique is an adaptation of what has been and is still used in various other research fields (e.g., volumetry of olfactory bulbs [19, 20], body parts and organs [21–23], etc.) and has proven to be a reproducible and reliable method. As expected, our evaluation of the reliability of the WDM proved to be positive with an excellent ICC (over 0.9), which proves the reliability of the WDM in measuring the volume of the full *ex vivo* human brain.

### 4.2 MRI manual and automatic segmentation volumetry

Our results showed that the MRI T2w manual, T2w automatic, and MP2RAGE INV1 automatic segmentation methods underestimated the total brain volume and that INV2 automatic segmentation overestimated it when compared with the gold standard of the WDM. This statistical difference may be due to the partial volume effect (PVE), commonly described as the loss of small tissue regions due to the limited resolution of the MRI scans [1]. Since one voxel can be composed of more than one tissue type, the resulting voxel intensity is an average of blurred tissue boundaries that is not characteristic of either underlying tissue [16]. Therefore, we can expect segmentation methods to underperform due to the PVE blurring the boundaries between different tissue types (e.g., all air-tissue or liquid-tissue interfaces).

A visual inspection of the segmentation masks produced on the T2w images enabled us to pinpoint the locations where the manual and automatic segmentations differ, with an underestimation of the manual masks, which are significantly smaller. We showed that the main differences occurred at the boundaries between the cerebral and cerebellar cortical matter and the surrounding CSF, as well as between the periventricular tissue and the intraventricular CSF. We hypothesize that a more inclusive threshold that segments more voxels around the boundaries between the cortex and CSF would bring the manual closer to the automatic segmented volumes, and both closer to the ground truth (i.e., the results generated by the gold standard WDM). It is also possible that the higher CSF-to-parenchyma contrast of the T2 image creates a higher PVE that may decrease the agreement across segmentation methods (manual vs. automatic) and across sequences. This is supported by the fact that even with a higher resolution than clinical standard acquisitions (0.5 mm^3^ against 1 mm^3^), the PVE problem persisted.

Another problem related to contrast between tissue types is when the contrast is decreased, for example between CSF and cortical GM, as is the case of the MP2RAGE INV2 sequence, which produced a misclassification of CSF voxels as GM voxels, falsely increasing the automatic INV2 volume in relation to the WDM.

However, we did not find any statistically significant differences between the WDM and MRI T1w automatically segmented brain volumes. This could possibly indicate that *ex vivo* T1w scans are less susceptible to PVE biases in the interface of tissue and CSF than other MRI sequences and that they more accurately represent the ground truth.

It is important to note that even though the statistical analysis did not find any significant differences between the T1w automatically segmented and WDM volumes, we still observed a mean difference of 16.45 cm^3^ between both volumetry methods. While this is not statistically relevant, it might be biologically relevant. Considering the mean human brain volume without the ventricles has previously been reported to be between 1090 and 1300 cm^3^ [24–27], the volume difference between both methods would represent about 1.3% to 1.5% of the total volume of the brain. Since we know from previous studies that the mean annual atrophy rate of the aging brain revolves around 0.2% to 0.5% [28–32] and can increase up to 2.4% in certain regions [28], our volume difference as it is observed when measuring the volume on MR images, even in T1w MRI scans where it was not statistically different from the WDM, needs to be taken into perspective. However, this discrepancy can be resolved by conserving the same MRI scanner, protocol, and sequence throughout any longitudinal study, as not to create method-related volume biases falsely passing for incorrect atrophy rates.

### 4.3 Application to an *in vivo* setting

Our validation method aimed to assess the accuracy in measuring brain volume of the MRI segmentation methods that are clinically available to be used for volumetric measures and research settings. Since we cannot extract the *in vivo* brains to measure their volume via a WDM, we had to think of an alternative solution that would allow us the closest estimation possible to *in vivo* brain volumes. We used the *ex vivo-in situ* perfusion method as described by Maranzano [10] to fix the brains directly inside the skull to allow us to scan the anatomical heads in a similar environment as *in vivo*, and to limit both the mechanical deformations of the tissue [12, 33] and the suboptimal quality that MRI scans may exhibit when they are fixed *ex situ* (i.e., artefacts caused by air bubbles remaining in the subarachnoid spaces around the brain, ventricles and blood vessels, degradation of superficial air-exposed layers, etc.) [10, 34, 35].

On the one hand, the *ex vivo-in situ* perfusion method offers the closest anatomical parallel to an *in vivo* environment. On the other hand, this means that we had to work with *ex vivo*, chemically fixed brain specimens. Indeed, they need to be fixed to preserve the cellular architecture and composition, and prevent decomposition, putrefaction and autolysis [36–39]. The chemicals used in the fixative solution alter the physicochemical characteristics of the tissue [33], which has an impact on the tissue properties (e.g., deformations causing anatomical and environmental differences, etc.) [8, 40–42] as well as on the MRI properties (e.g., modification of relaxation values and/or contrasts) [19, 42–46]. This changes the contrasts between GM-WM-CSF of the MRI images, which might in turn affect the quality of the generated automatic segmentation, since the SynthSeg software was not designed specifically for dealing with *ex vivo* specimens.

However, the T2w sequences of our *ex-vivo-in situ* brains are extremely close to an *in vivo* T2w scan (when visually assessed by an expert; J.M), so we were not overly concerned with the impact of the fixative on the tissue from an anatomical standpoint (because we did not see a particular deformation in any brain area) nor from a voxel-intensity standpoint (because we observed a similar GM-to-WM-to-CSF contrast as in *in vivo* T2w scans). Additionally, as mentioned before, dealing with the *ex vivo* specimens was advantageous since we could establish the correlation with the gold standard method, and increase the MRI quality by using protocols that are not time-sensitive, motion-free and at a higher resolution than standard *in vivo* scans [1, 8, 19, 40–42, 44–46].

More importantly, the closest MRI automatic volumes to the ground truth (i.e., the WDM volumes) were obtained using a T1w sequence, which happens to be the most frequently used sequence *in vivo* for brain volumetric analysis. While we are aware that our *ex vivo* T1w images exhibit different GM-to-WM contrasts than the *in vivo* scans, our *ex vivo* T1w images conserve the CSF-to-GM contrast that is typical of *in vivo* T1w (i.e., hypointense CSF in relation to a more intense cortical GM). We surmise that it gives a better depiction of the CSF-GM boundary that explains the superior results of automatic volumes using T1w images.

Unfortunately, the UNI MP2RAGE sequence, which is also commonly used in clinical settings, did not generate reliable volumes, although the problems stemmed from the background noise causing a voxel tissue type misclassification of certain anatomical structures (mainly in the brainstem). For this sequence, additional post-processing steps of the automatic output will be necessary if it is to be used in automatic brain volumetric analyses.

### 4.4 Limitations

Our study consists of a small sample size (N=7) composed of the available brains from the UQTR’s Anatomy department, so the conclusion of this work might not be applicable for specimens fixed with other chemical solutions. To that end, we are planning to expand our sample size as more brain specimens become available to us. We will also measure the volumes in heads fixed with a different fixative solution.

At this stage, we have not yet perfected the post-processing steps that would be required to improve the automatic volumes on the UNI MP2RAGE images. Future work will determine if the cost-benefit analysis of finding and applying the right corrections is worth the investment of time and resources to make this sequence usable or if continuing to work with the T1w and T2w images is sufficient.

Finally, the software use in this work (SynthSeg) generates outputs at 1mm^3^ resolution regardless of the input voxel size, which adds to the PVE in the measured volume after resampling the segmentation labels back to their native resolution when using sub-millimeter brain images.

### 4.5 Conclusion

We propose herein that the MRI automatic segmentation method using SynthSeg can be used to accurately determine the brain volume on *ex vivo* T1-weighted MRI scans, which proved to generate volumes that were not statistically different from the volumes measured with the gold standard WDM. Furthermore, the contrast similarity of our *ex vivo* sequences to *in vivo* sequences (specifically of T2w images, where *in vivo* and *ex vivo* are visually indistinguishable) suggests that brain volumes in clinical studies settings are most probably sufficiently accurate too, even if some sequences may under or overestimate the ground truth, which is not a problem if the same scanners are used.

## Author contribution

DB and JM contributed to the conception and design of the study. AGL performed the experimental protocols, organized the database and performed the statistical analysis. WAG, EMF, LGS, AN, MD, and JM performed a part of the experimental protocols. AGL wrote the article. JM wrote sections of the article and contributed to manuscript revision. All authors read, revised, and approved the submitted version.

## Acknowledgements

First, we would like to thank the body donors and their families for their generosity and for making this research possible. We also acknowledge all our funding resources and our anatomy laboratory personnel at the Université du Québec à Trois-Rivières for their contribution (Johanne Pellerin, Marie-Eve Lemire, Sophie Plante, Sonia Gauthier, and Kevin Desaulniers). Lastly, we would like to thank the staff at the Cerebral Imaging Centre (The Douglas Research Centre) for their help in acquiring the images.

